# Female cortical cellular mosaicism underlies shared MeCP2 and PCB impacted gene pathways

**DOI:** 10.1101/2025.05.28.655836

**Authors:** Osman Sharifi, Kari E. Neier, Anthony Valenzuela, Christina G. Torres, Ian Korf, Pamela J. Lein, Dag H. Yasui, Janine M. LaSalle

## Abstract

Etiologies of neurodevelopmental disorders involve genes and environment however their interactions are understudied. Rett Syndrome (RTT) is an X-linked, dominant neurodevelopmental disorder caused by mutations in *MECP2*, encoding the epigenetic regulator methyl CpG binding protein. Epigenetic features of *MECP2* expression due to X-linked cellular mosaicism and the variability in severity and timing of progression in RTT suggest interaction with environmental neurotoxicants such as lipophilic polychlorinated biphenyls (PCBs). To understand shared mechanisms, we exposed WT and *Mecp2e1*^-/+^ female mice to a human-relevant PCB mixture and dose, then performed single-nucleus 5’ RNA-seq from cortex. We identified significant overlap in dysregulated genes and 71 shared pathways between the effects of PCB exposure and MeCP2 mutation, and co-mitigation of their transcriptional impacts. PCBs influenced the non-cell-autonomous transcriptional effects of MeCP2 mutations in wild-type-expressing neurons within the mosaic mutant female cortex in both mouse and human, suggesting that the interactions predominantly involve homeostatic gene networks.

## Introduction

In female mammals, X-chromosome inactivation (XCI) is random, affecting either the maternal or paternal allele in each cell. Once established, this creates a cellular mosaicism throughout the body, unless there is selective loss of one pattern^1^. Rett Syndrome (RTT) is an X-linked dominant neurodevelopmental disorder that primarily affects females and is caused by mutations in *MECP2*^2^. This gene encodes Methyl CpG binding protein 2 (MeCP2), a critical epigenetic regulator that modulates gene expression by binding to methylated DNA and acting as either a repressor or activator of transcription^3–5^. RTT infants are born seemingly healthy but experience a significant regression in cognitive and motor skills during late infancy, followed by several subsequent stages of regression through young adulthood, highlighting the importance of MeCP2 for the maintenance of normal neurological function^6^.

The progression and severity of RTT are highly variable, suggesting that the disease pathogenesis may be influenced by additional factors beyond genetic mutations alone. Using single nucleus RNA sequencing (snRNA-seq 5’), we recently demonstrated a dynamic, non-cell autonomous effect in wild-type expressing cells within the cortex of a construct-relevant model of RTT that corresponded with disease progression^7^. This variability and cellular mosaicism raises the possibility that *MECP2* mutations might interact with environmental factors, further complicating the disease’s clinical presentation^8^. Environmental toxicants such as persistent organic pollutants (POPs) have emerged as potential contributors to neurodevelopmental disorders (NDDs)^9,10^. Among these, polychlorinated biphenyls (PCBs) are notable for their widespread environmental presence and their association with adverse neurodevelopmental outcomes^11^. PCB 95, in particular, has been shown to promote dendrite growth and synaptogenesis^12,13^. We previously examined the differentially methylated regions (DMR) of fetal brain and placenta of mice exposed to PCBs and revealed a significant overlap between MeCP2 regulated genes and differentially methylated genes resulting from prenatal PCB exposure^8^.

PCBs have been detected in the brains of individuals with various NDDs, including RTT^14^. Research has demonstrated that exposure to PCB 95 can alter the DNA methylation patterns of over 1,000 genes in cell lines, suggesting a potential interaction between genetic defects and environmental exposures in the disruption of chromatin epigenetics^15^. Furthermore, the Markers of Autism Risk Learning Early Signs (MARBLES) study has measured PCB congeners in mothers at increased likelihood for autism spectrum disorders (ASD), highlighting the relevance of PCB exposure in developmental neurotoxicology^16^. Exposing mice to a mixture of the PCB congeners identified in the MARBLES study led to an increase in dendritic branching of cortical neurons^17^ and significant deficits in sociability and ultrasonic vocalizations, as well as enhanced repetitive behavior specifically in males at the lowest dose tested (0.1 mg/kg/d)^18^.

To explore the intersection of genetic and environmental factors in RTT female X chromosome mosaicism, we utilized a construct-relevant mouse model of RTT and performed snRNA-seq 5’ on cortical cell types from adult female mice exposed to the MARBLES PCB mix. We demonstrate a highly significant overlap in dysregulated genes and molecular pathways and between the effects of PCB exposure and *Mecp2* mutations. Interestingly, PCBs preferentially impact the non-cell-autonomous transcription effects of *Mecp2* mutation in wild-type-expressing neurons within *Mecp2* mosaic mutant female cortex, suggesting the interaction is predominantly with homeostatic gene networks. By establishing a baseline for single-cell gene expression in PCB-exposed mice, our study aims to elucidate the complex interplay between genetic and environmental factors in RTT and related neurodevelopmental disorders, providing insights into how PCB exposures may modulate disease progression and severity.

## Results

### Experimental design to test transcriptional dysregulation in PCB-exposed WT and *Mecp2e1* heterozygous female cortical cell types

To identify cell type and PCB exposure specific transcriptional differences in *Mecp2e1* deficient mouse cortex, single nuclei RNA sequencing (sn-RNA seq 5’) analysis was performed to capture the translational stop site mutation at the 5’ translational start site of the *Mecp2e1* isoform^19^. Late disease stage *Mecp2e1^-^*^/+^ (∼16 weeks of age) heterozygous (HET) females were chosen and compared to sex-matched wild-type *Mecp2e1^+^*^/+^ (WT) littermates (**Figure 1A**). Female mice of each genotype were orally exposed to either the PCB MARBLES mix or vehicle control daily for ∼6 weeks for a total of 16 animals (4/group) (**Figure 1A**). To elucidate the cellular landscape within the cortices of WT and HET mice, we applied unsupervised clustering analysis to classify cells based on their gene expression profiles and labeled cell types based on the Allen Brain Atlas cortex reference^20^. We identified 12 distinct cell types within the cortical tissue (**Figure 1B**). Cell type identification and labeling was validated based on marker genes present in each cell type (**Figure 1C**). Each marker was selected based on its known association with specific cell types and validated the accuracy of our clustering results. To assess the influence of PCB exposure on cell type clustering, we compared the clustering results of cells treated with the PCBs to those treated with a vehicle control (**Figure 1D**). Cells from both treatment conditions were clustered mostly together, and the resulting cell type distributions were analyzed for differentially expressed genes (DEGs).

**Figure 1.**
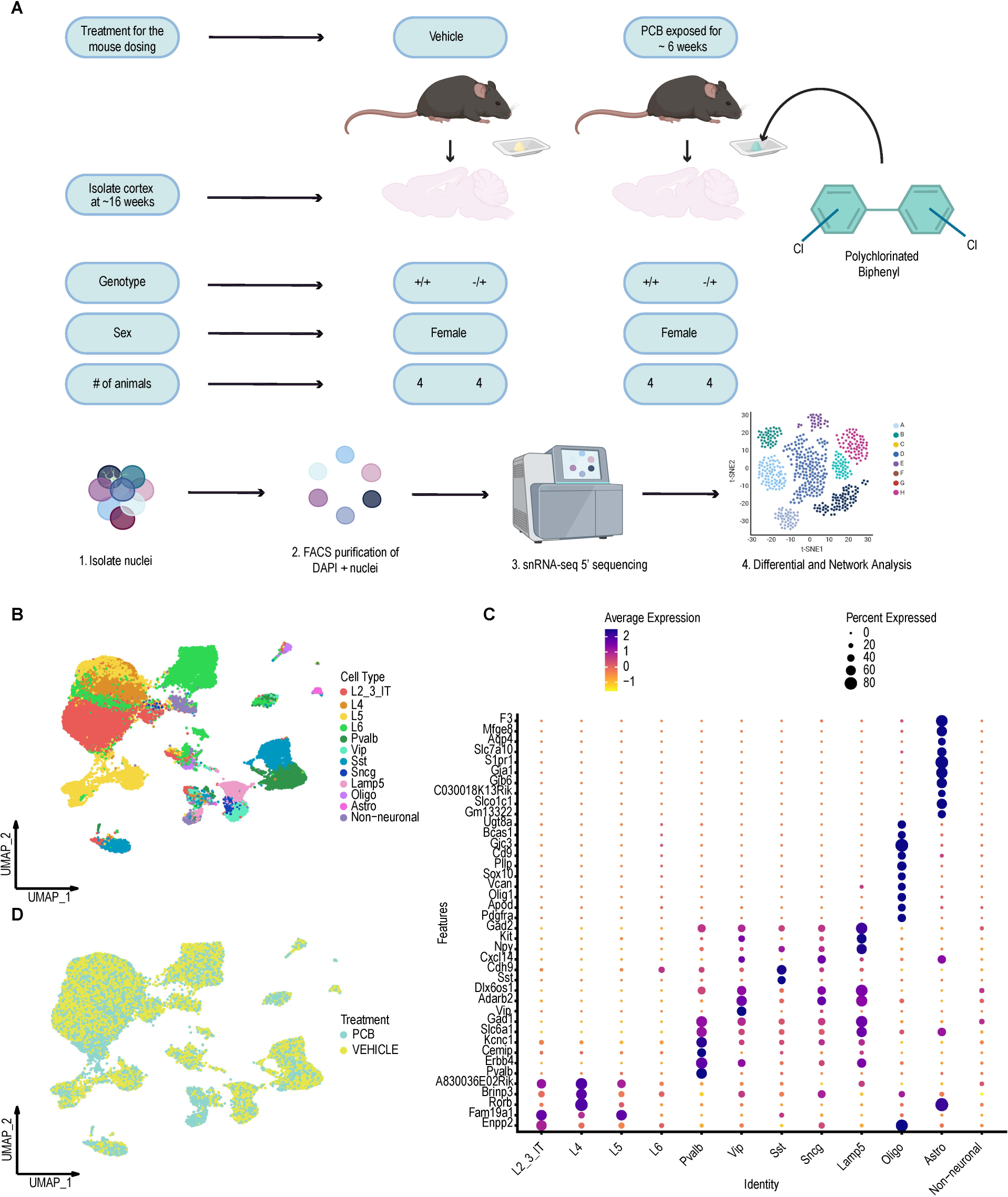
: Single nuclear transcriptome study design to investigate effects of *Mecp2* mutation and PCB exposure. **A.** Schematic showing cortical samples from PCB or vehicle treatment groups of two genotypes (*Mecp2e1^+/+^* WT or *Mecp2e1^-/+^* HET) for snRNA-seq 5’. **B.** UMAP unsupervised clustering of cell types identified in cortex from all samples. **C.** Dot plot of the top marker genes for each cell type identified. **D.** UMAP plot of cell cluster by treatment group.

### Cell type stratified differential gene expression (DEG) analysis of cell types in PCB-treated versus vehicle-treated females of both *Mecp2e1* genotypes

To further explore the impact of PCB exposure on gene expression within distinct cell types, we performed a differential expression analysis comparing vehicle-treated and PCB-treated conditions in both WT and HET mice. This analysis aimed to identify changes in gene expression associated with PCB exposure and evaluate potential interaction effects of PCB treatment in the context of RTT HET genotype (**Figure 2**). To ensure DEGs were not driven by changes in cellular proportions, we conducted an analysis of cell type proportions in both WT and RTT cortices (**Supplemental Figure 1, Supplemental Table 1**) and did not observe significant changes consistent with a previous study^21^. We first compared cell type DEGs between WT and HET mice within the vehicle-treated and PCB-treated backgrounds. This comparison aimed to discern how PCB exposure interacts with genetic differences between WT and HET mice. The variability in number of DEGs and expression patterns across different cell types underscores the complexity of PCB’s effects, which may vary depending on the specific cellular context and the nature of the gene expression changes (**Figure 2A**). The highest number of DEGs were observed between WT and HET cortical cells with vehicle treatment. Conversely, PCB treated HET mice showed a lower number of DEGs compared to vehicle treated HET cortical cells (**Figure 2A**). These findings highlight a potential modulatory role of PCB in the context of HET, where PCB treatment may normalize or attenuate some of the transcriptional disruptions caused by the *Mecp2e1* mutations. These DEGs are primarily associated with neuronal function, brain development, signaling pathways, and metabolism (**Figure 2B-C**). Many encode proteins in the nervous system, including neurotransmitter regulation such as *Gad1* and *Slc1a2*, calcium signaling (*Calm2* and *Hpca*), and transcription factors involved in development (*Zbtb20* and *Meis2*) (**Figure 2B-C**). Comparisons of PCB-treated vs. Vehicle and WT cortices vs. RTT cortices both contain genes involved in neuronal signaling and synaptic function (*Syt1, Nrgn, Calm1/Calm2, Cnr1, Slc1a2, Pcp4*), suggesting a dysregulation of neuronal communication.

**Figure 2:**
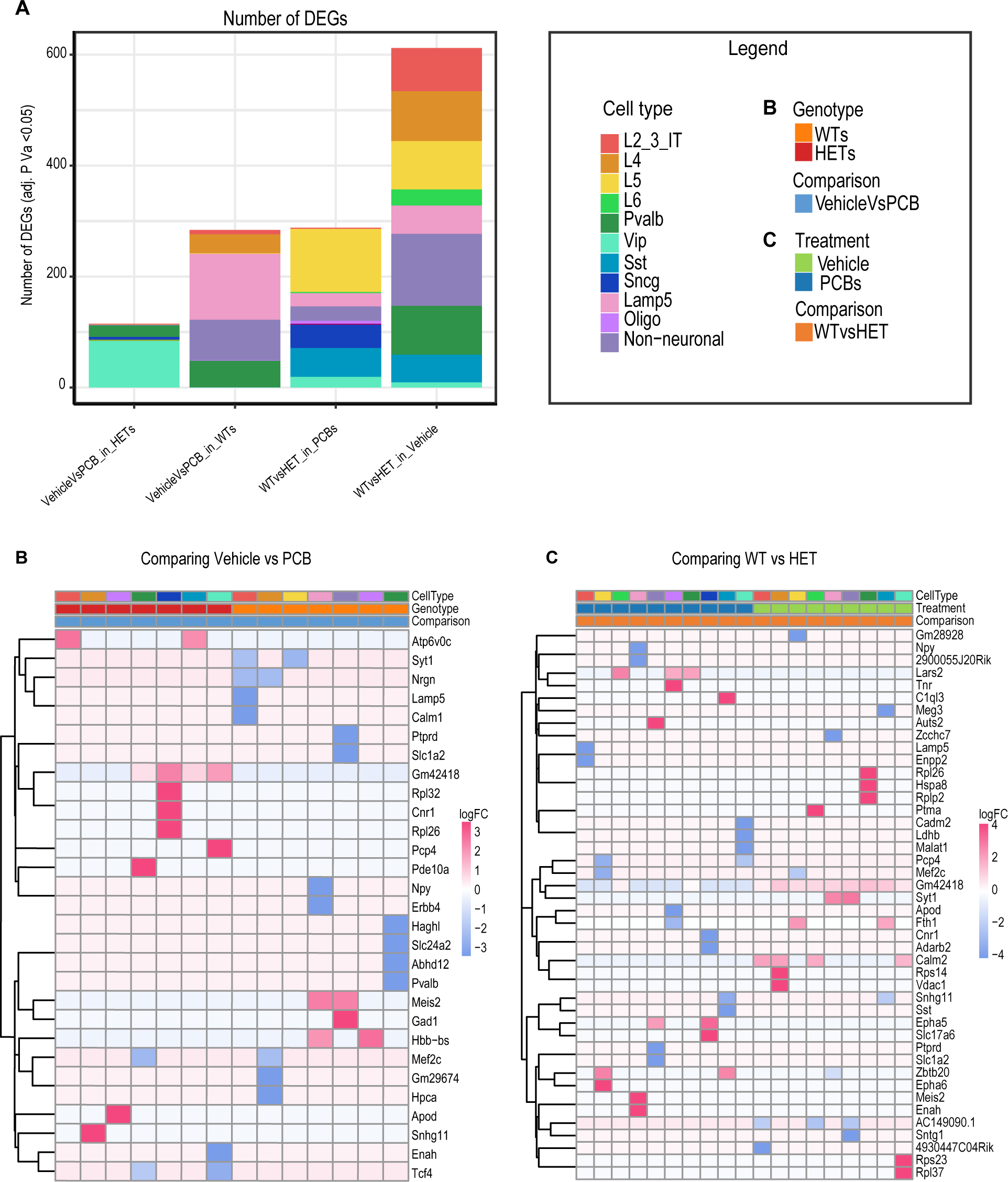
Interaction between *Mecp2* genotype and PCB exposure in frequency and cell type distribution of differentially expressed genes (DEG). **A.** Bar graph showing the significant number of DEGs for each experiment. **B, C.** Heatmaps showing the top DEGs by log-fold change (adjusted p-value ≤ 0.05) per cell type for comparing Vehicle vs PCB and comparing WT vs Mecp2e1^-/+^ HET cortices respectively.

### Commonly altered KEGG pathways are observed across experimental conditions

To understand how PCB exposure affects cellular pathways, we analyzed dysregulated KEGG pathways across four pairwise comparisons of cell type DEGs: WT vs HET in vehicle, WT vs HET in PCB, vehicle vs PCB in WT, and vehicle vs PCB in HET. This analysis aimed to identify common and unique pathway alterations associated with the interaction between PCB exposure and heterozygous *Mecp2e1* mutation. While the total number of KEGG pathways did not vary by comparison group, non-neuronal cell types showed a reduction in KEGG terms specifically in the genotype comparison within PCB treated animals (**Figure 3A**). A closer look at the dysregulated pathways reveals that the majority of these KEGG pathways are highly similar across all four group comparisons, with 71 overlapping pathways (**Figure 3B**). This extensive overlap indicates that both PCB exposure and *Mecp2e1* genotype converge on a core set of dysregulated gene pathways. Across four experimental conditions, these overlapping dysregulated gene pathways were more impacted in neuronal than glial cells (**Figure 3C, Supplemental Table 2**). Further, the KEGG pathways that were dysregulated across conditions and cell types include those related to neuronal and synaptic function such as GABAergic synapse, glutamatergic synapse, synaptic vesicle cycle, long-term potentiation (LTP), long-term depression (LTD), circadian rhythm and circadian entrainment, amphetamine addiction, morphine addiction, and nicotine addiction (**Figure 3C**). These neurologically relevant pathways underscore the potential interaction of PCB exposure and a neurodevelopmental disorder mutation on key aspects of neuronal function and regulation. In addition, many of the dysregulated paths are related to neurogenerative diseases such as Parkinson’s disease and Alzheimer’ disease, which is consistent with previous studies^22,23^.

**Figure 3:**
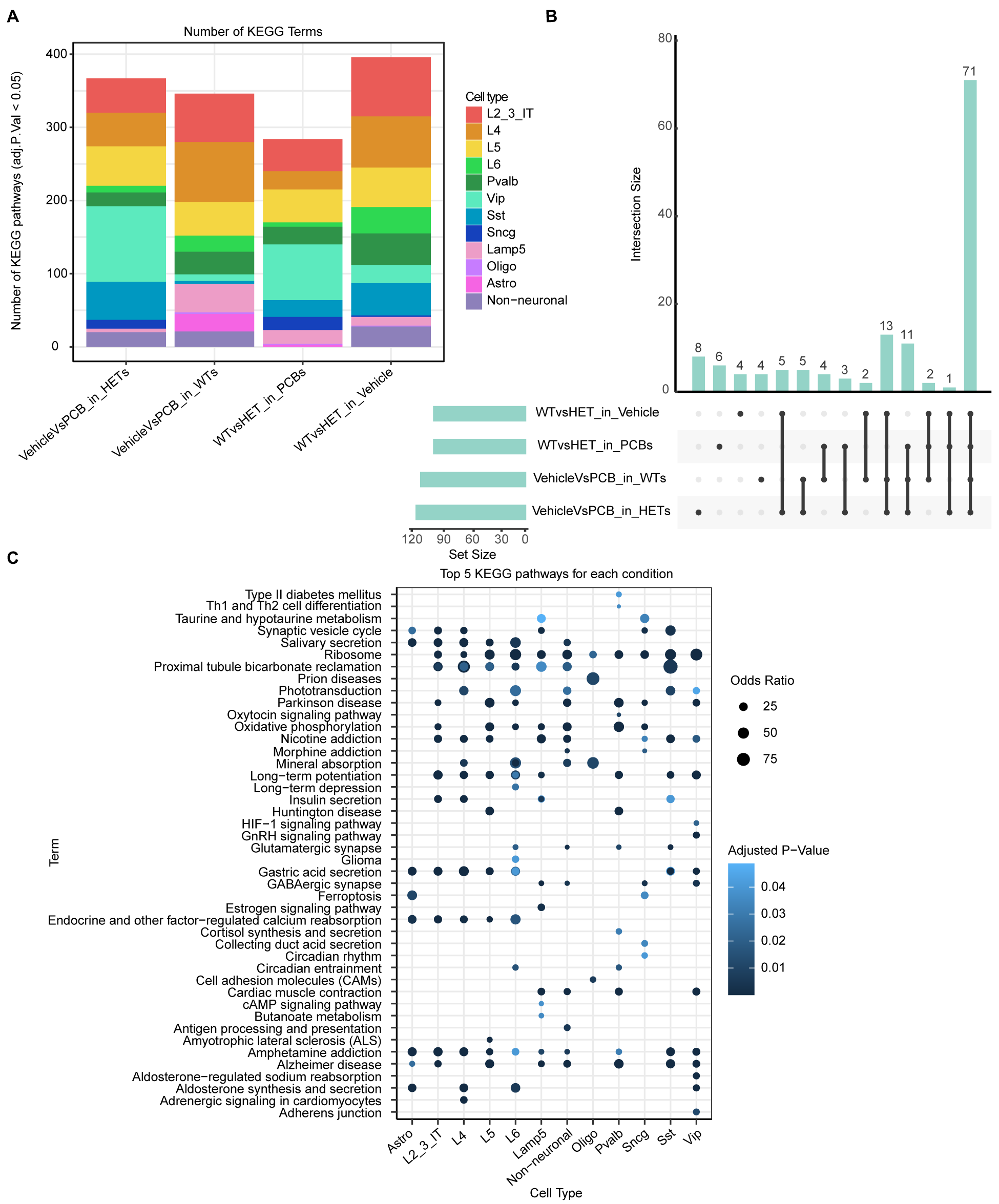
**KEGG pathway analysis shows convergent biological pathways between *Mecp2* genotype and PCB exposure**. **A**. Bar graphs showing the number of significant dysregulated KEGG pathways per cell type. **B**. Upset plot showing the number of significant pathways overlapping across the 4 experimental pairwise comparisons. **C**. Dot plot showing the top overlapping pathways per cell type.

### Investigating PCB effects on *Mecp2e1* heterozygous mosaic cortices and cell non-autonomous interactions

To accurately distinguish between wild-type (WT) and mutant *Mecp2*-expressing cells within a mosaic female brain, we developed a bioinformatic pipeline (**Supplemental Figure 2**). This pipeline integrates preprocessing, feature selection, dimensionality reduction, clustering, and classification to achieve precise segregation of WT and mutant *Mecp2*-expressing cells in the mosaic brain in the context of our gene X environment experimental design.

To investigate how PCB exposure affects WT-*Mecp2* versus mutant-expressing cells within female mosaic cortical tissue, we first parsed *Mecp2* transcript genotypes in all cells (**Figure 4A-D**). As expected, no mutant cells were observed in WT cortices (**Figure 4D**), and female heterozygous cortical cells were a roughly equal mixture of WT- and mutant-expressing (**Figure 4A, C**).

**Figure 4:**
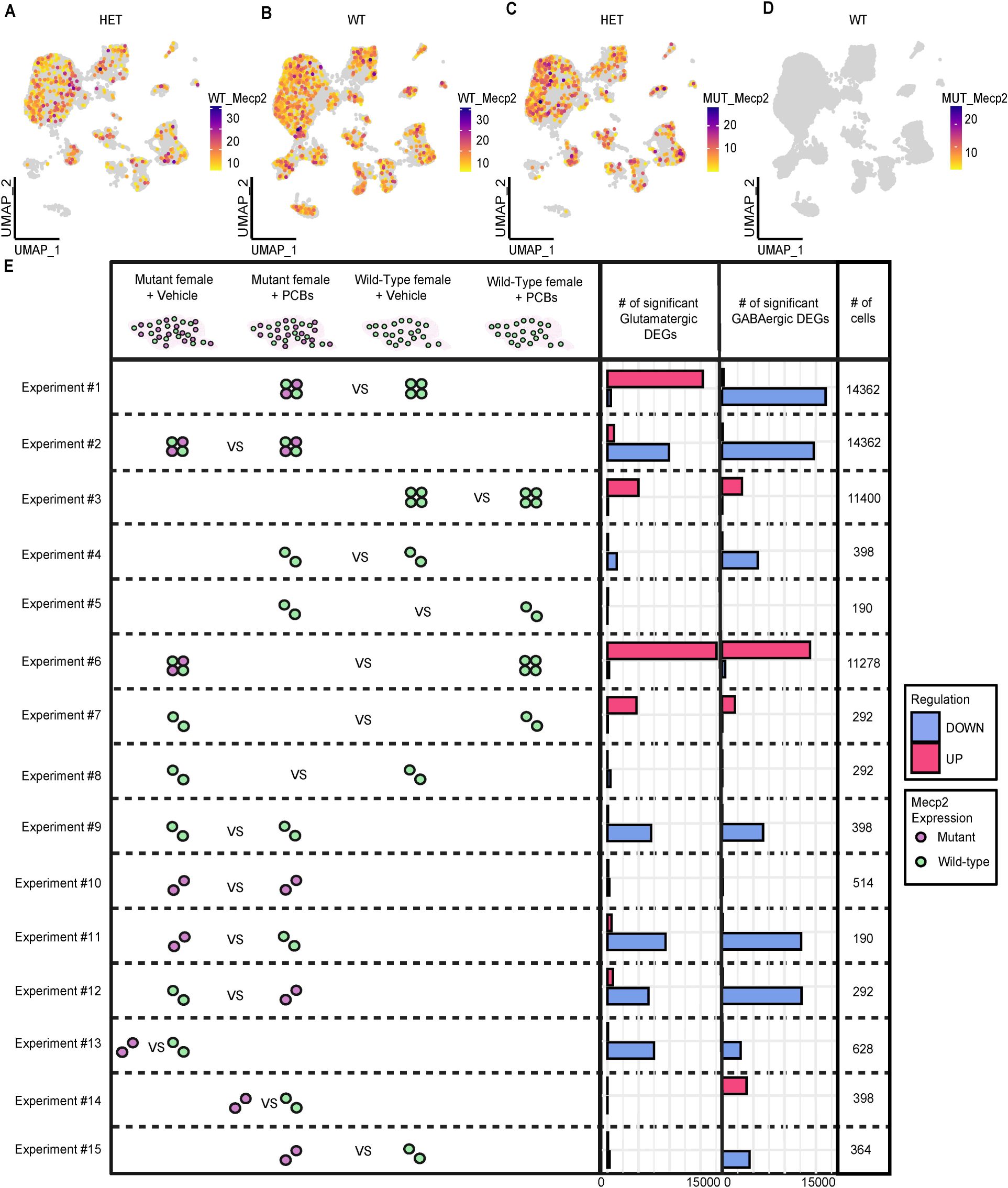
Mosaic cellular parsing reveals PCB effects on non-cell-autonomous transcriptional dysregulation in RTT mouse model. A, B, C, **D**. UMAP plots showing WT_*Mecp2* and MUT_*Mecp2* expression in WT and *Mecp2e1* mutant heterozygous cortices. **E.** DEG experiments comparing WT (green) or mutant (purple) single cells across sample type in Glutamatergic and GABAergic cells15 experimental comparisons (y-axis) for DEGs (bar graphs; red, up; blue, down for right group) revealed opposite PCB effects on Glutamatergic vs GABAergic neurons in bulk HET (#1) and opposite effects of mutant-expression within HET vs WT cortical GABAergic neurons (#14-15). No mutant-expressing cells were detected in WT cortex, as expected. Cells were down-sampled to the lowest number of cells per pair, n=3 mice/group

To reduce the impact of lower cell counts on DEG calling following parsing, we further grouped the *Mecp2* expressing cells into neuronal categories of GABAergic and glutamatergic neurons to gain insight into cell non-autonomous effects. The number of cells analyzed are given for each comparison after down-sampling so that the pairwise comparisons were equal (right column **Figure 4E, Supplemental Table 3**). Non-neuronal cells were not sufficiently abundant to allow for DEG analyses after parsing. To evaluate the effects of *Mecp2* genotype and PCB exposure on neuronal populations, we conducted 15 distinct differential expression (DEG) experiments, highlighting the impact of PCB exposure on each *Mecp2* genotype in each of these neuronal subtypes (**Figure 4E**). Experiments #1, 2, 3, and 6 are from non-parsed (4 dot) Glutamatergic or GABAergic neurons for comparison to parsed (2 dot) neurons in remaining experiments that were either WT-expressing (green) or mutant-expressing (purple). Interestingly, a comparison of PCB effects in WT-expressing cells compared to mutant-expressing cells in HET (experiments 9 and 10) showed a higher number of DEGs in WT-expressing neurons than mutant-expressing neurons of both types. Remarkably, the lowest numbers of DEGs were seen when comparing WT-expressing neurons across both genotype and treatment groups (experiments #5 and 8) or mutant-expressing neurons with or without PCBs (experiment #10). This indicates that PCB exposure predominantly affects *Mecp2* WT-expressing neurons, suggesting that PCB interaction effects within RTT cortices are predominantly acting on the homeostatic gene networks dysregulated in WT-expressing neurons. Comparisons of non-parsed neurons by mouse *Mecp2* genotype with or without PCB (experiments #1 and 6, respectively) showed that the number of and direction of DEGs were different between excitatory and inhibitory neurons only in the presence of PCBs. Collectively, these results suggest that PCB treatment interacts with and could mitigate some of the detrimental non-cell-autonomous effects caused by *Mecp2* mutation (**Supplemental Table 4**). The differences observed in DEG counts between *Mecp2e1* WT- and mutant-expressing cells, and the impact of PCB treatment on these cells, highlight the complex interactions within the mosaic brain environment. PCB exposure appears to interact with *Mecp2e1* mutation by transcriptionally dysregulating shared homeostatic pathways regulating the broader cellular milieu, demonstrating a nuanced interaction between a genetic and environmental factor.

### Network analysis using hdWGCNA associates PCB and *Mecp2e1* genotype effects with RTT mouse model phenotypes

To further explore the effects of *Mecp2e1* mutations and PCB exposure on RTT phenotypes, we employed high dimensional Weighted Gene Co-Expression Network Analysis (hdWGCNA). This approach is relevant for assessing gene networks within snRNA-seq data and allowed us to analyze associations of experimental variables (treatment, genotype, exposure duration, and pregnancy) and measured phenotype (body weight at time of sacrifice at ∼16 weeks) with gene co-expression networks. hdWGCNA analysis results show that the gene network is organized into seven distinct modules (**Figure 5A**). Each module represents a cluster of co-expressed genes with common functional characteristics. Hub genes within each module are highlighted, serving as central connectors that are crucial for the module’s function and overall network stability (**Figure 5A**). The genes from these distinct modules were used to perform KEGG pathway enrichment analysis. The top KEGG pathways for each module reflect the biological processes and pathways most prominently affected by PCB exposure and/or *Mecp2e1* genotype (**Figure 5B**). These pathways include many neurological disease related terms such as Parkinson’s disease, Alzheimer’s disease, calcium signaling and long-term potentiation which provide insights into the functional impact of these factors on gene networks (**Figure 5B**). To confirm that the gene networks are not influenced by cell type-specific gene markers, we performed an overlap analysis between cell type gene markers and the identified gene modules (**Figure 5C**). Additionally, we examined the correlation of gene expression in each cell type with various experimental variables, including pregnancy, weight at sacrifice, exposure duration, PCB treatment, and genotype (**Supplemental Figure 3**). Among the different cell types analyzed, the Sst inhibitory neurons showed the strongest correlations with changes in gene expression linked to genotype and phenotypic traits, suggesting that *Mecp2e1* genotype and PCB exposures interact within shared excitatory/inhibitory balance and homeostatic transcriptional networks in the female adult cortex.

**Figure 5:**
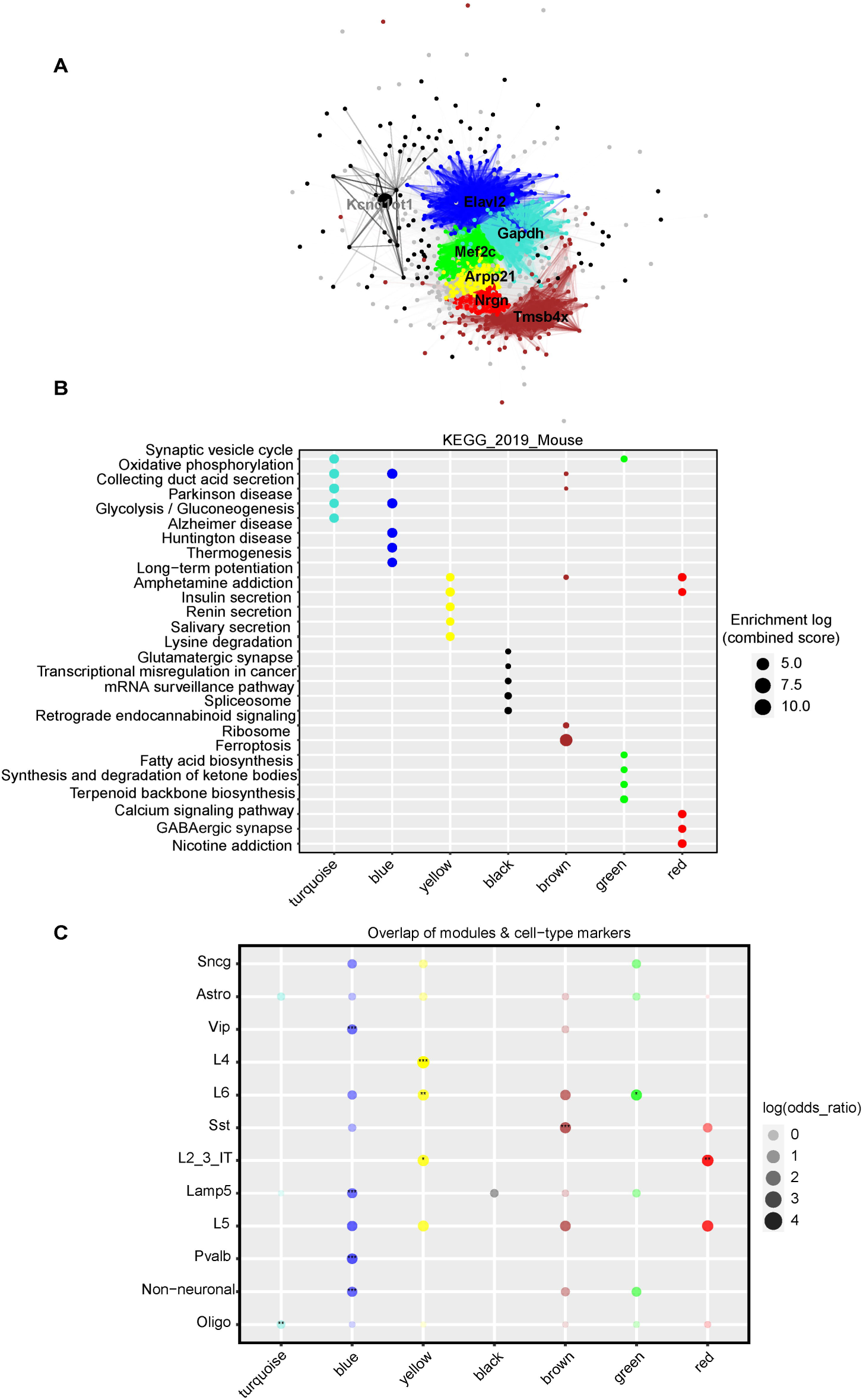
hdWGCNA analysis reveals correlated transcriptional networks impacted by *Mecp2e1* mutation and PCBs. **A**. Weighted gene network showing 7 modules and their corresponding hub genes. **B**. Dot plot showing the top 5 KEGG pathways for each module. **C**. Dot plot showing the overlap of cell type markers and gene modules.

### Comparative analysis of mouse and human Rett cortex PCB-associated DEGs and enriched pathways in GABAergic, Glutamatergic, and Non-Neuronal Cells

To evaluate the relevance of our findings in a mouse model to human Rett syndrome, we compared cell type-specific DEGs associated with PCB exposures between human female Rett cortex samples with measured PCB levels^7,14^ and mouse female HET cortex samples. **Figure 6** illustrates this comparative analysis, highlighting conserved and divergent molecular signatures across species.

**Figure 6:**
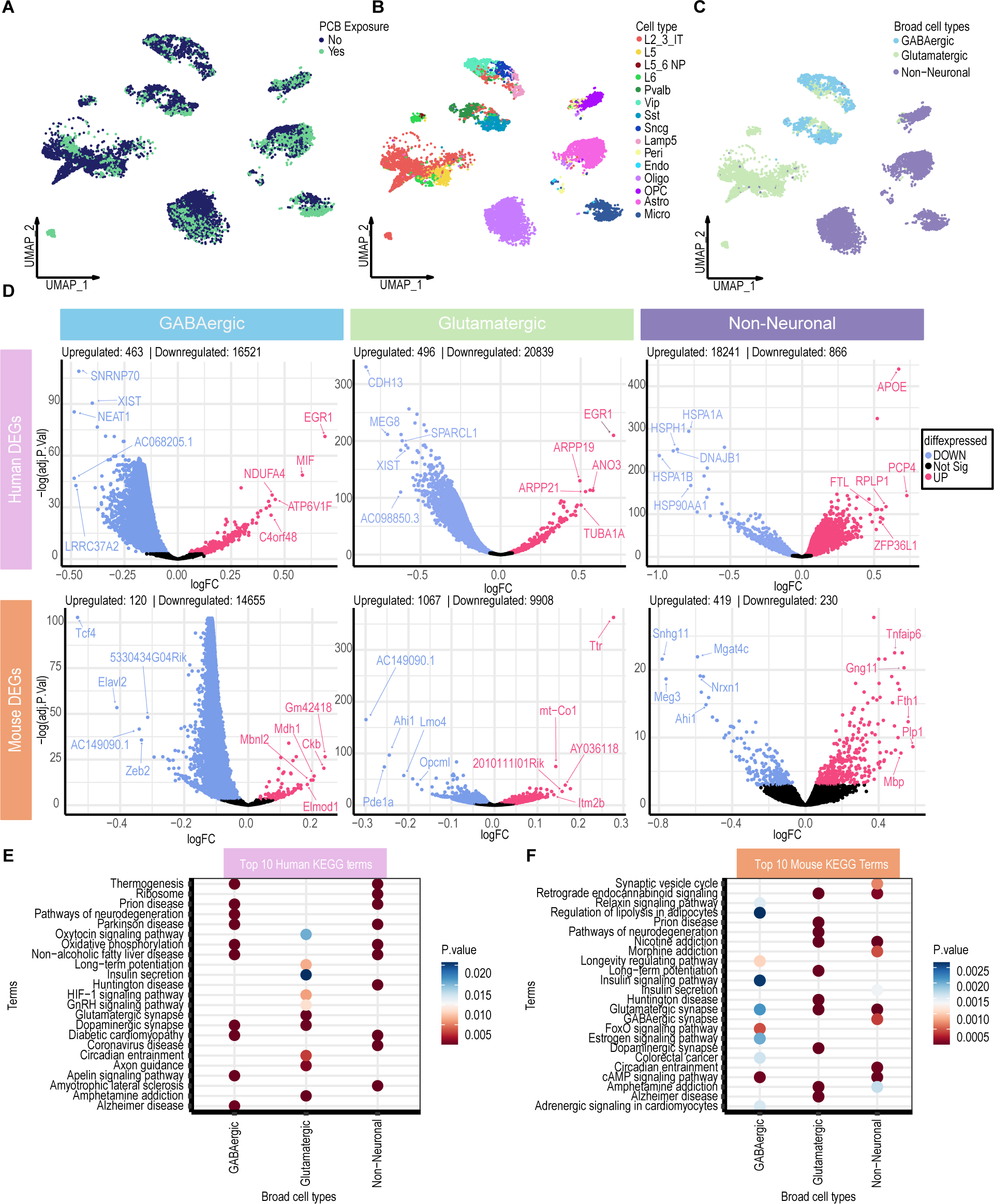
Comparative analysis of mouse and human Rett cortex PCB-associated DEGs in GABAergic, Glutamatergic, and Non-Neuronal Cells. **A**. UMAP plot of PCB and non-PCB exposed cell clustering. **B**. UMAP plot of unsupervised cell type clustering. **C**. UMAP plot of cell types grouped in two three broad cell types. **D**. Volcano plots of human and mouse DEGs in the three broad cell types. Significant DEGs are in Blue and Magenta. **E, F**. Dot plots of top 10 human and mouse KEGG terms respectively.

UMAP projection of human RTT brains, stratified by PCB exposure status (Yes means any PCB congener measured above level of detection), revealed similar cell clustering associated with exposure (**Figure 6A**). Human cell types were identified and annotated based on the Allen Brain Human Cortex dataset^24^ (**Figure 6B**). For a cross-species comparative analysis, these cell types were grouped into three major categories: GABAergic neurons, glutamatergic neurons, and non-neuronal cells (**Figure 6C**). DEGs comparing PCB-exposed and non-exposed conditions for each broad cell type in both human and mouse datasets were identified (**Figure 6D**). Strikingly, GABAergic and glutamatergic neurons in RTT cortices of both species exhibited a predominance of downregulated PCB-associated DEGs, suggesting conserved mechanisms of PCB associated transcriptional repression in these neuronal populations. In contrast, non-neuronal cells in both human and mouse show a trend of upregulated DEGs, indicating a shared activation of transcriptional programs in these cell types following PCB exposure (**Figure 6D**). The top KEGG pathways enriched in human and mouse datasets were identified (**Figure 6E, F, respectively**). Both species exhibit dysregulation in pathways related to synaptic function, neurodegeneration, and metabolic regulation. Notably, conserved pathways include synaptic signaling such as GABAergic synapse and glutamatergic synapse and retrograde endocannabinoid signaling in addition to metabolic and addiction pathways (**Figure 6E, F**). Interestingly, the female-specific noncoding *XIST* gene was a top PCB-associated downregulated DEG in human that was also overlapping with mouse. A highly significant interaction between *Mecp2e1* genotype and PCB exposure was observed for *Xist* levels, with cell type-specific differences (**Supplemental Figure 4**). These findings underscore the translational relevance of our mouse model to human Rett syndrome, demonstrating conserved molecular and cellular responses to PCB exposure in *Mecp2* mosaic female cortex across species. The shared dysregulation of key pathways involved in synaptic function, neurodegeneration, and metabolic homeostasis highlights potential mechanistic links between an environmental exposure and Rett syndrome pathology.

We further explored the conservation of molecular responses between mouse and human Rett cortex by analyzing the overlap of significant DEGs and KEGG pathways across GABAergic, glutamatergic, and non-neuronal cell types (**Figure 7**). The analysis of significant DEGs between mouse and human datasets reveals that the greatest overlap occurs in GABAergic cells (7931 DEGs), followed by glutamatergic cells (6344 DEGs), and lastly, non-neuronal cells (453 DEGs) (**Figure 7A**). This pattern suggests that similar genes are dysregulated in these neuronal populations across species, highlighting conserved transcriptional responses to PCB exposure within Rett cortex. The overlap of significant KEGG terms between mouse and human datasets across the three broad cell types were consistent with the DEG overlap. The greatest convergence of KEGG terms is observed in GABAergic cells, underscoring the shared molecular pathways affected in this cell type (**Figure 7B**). Notably, only one KEGG term, oxytocin signaling pathway, is common to both species and all three cell types, emphasizing oxytocin’s potential role as a conserved regulatory network of inter-cellular homeostatic responses in Rett syndrome.

**Figure 7:**
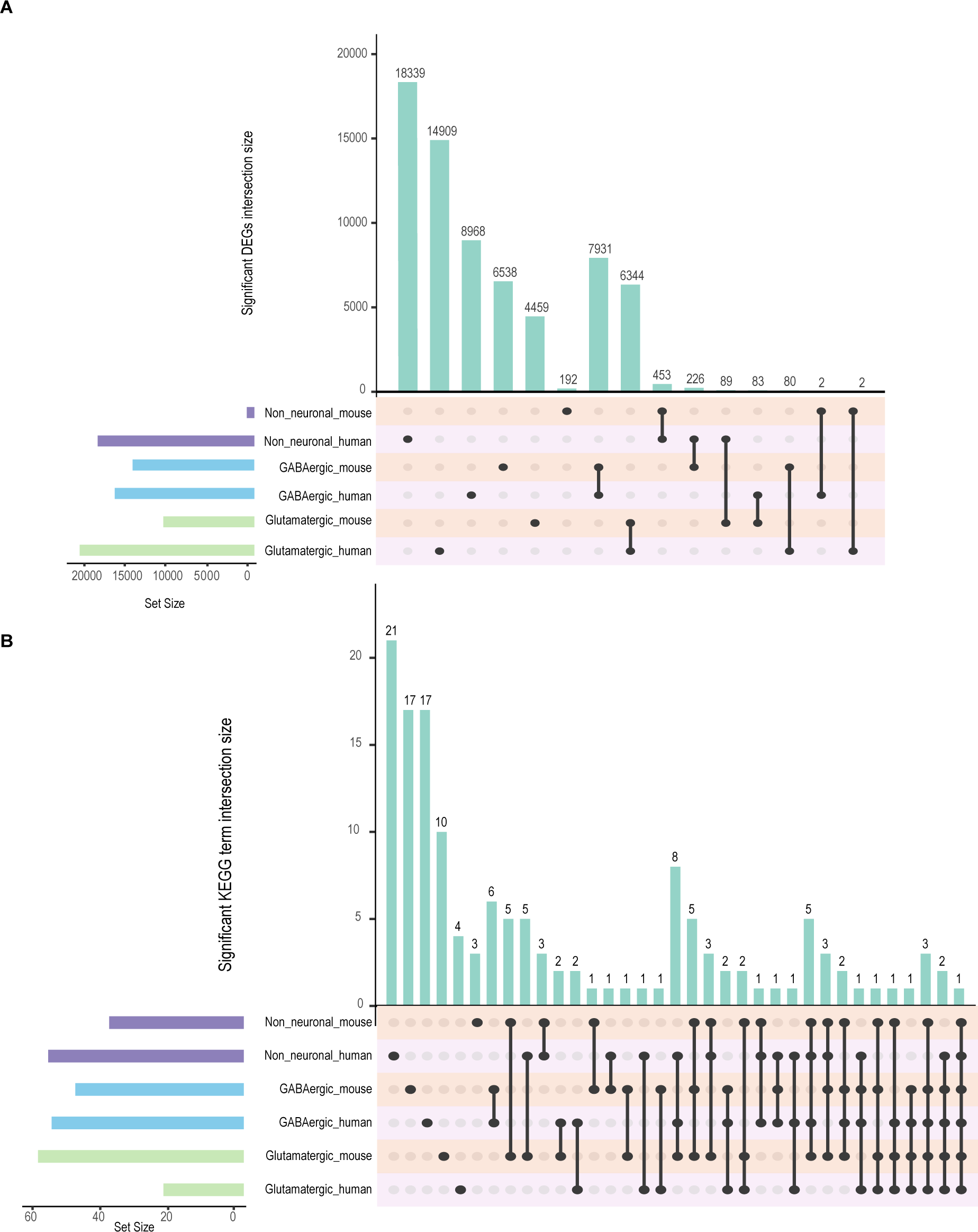
Overlap of mouse and human Rett cortex PCB-associated DEGs and KEGG terms in GABAergic, Glutamatergic, and Non-Neuronal Cells. A, **B**. Upset plots of significant (adjusted p-value ≤ 0.05) DEGs and KEGG terms across human and mouse broad cell types respectively.

These findings reinforce the translational relevance of the mouse model to human RTT, demonstrating conserved gene and pathway alterations, particularly affecting GABAergic and glutamatergic neurons. The identification of oxytocin signaling as a shared pathway across species and cell types suggests its potential importance in the pathophysiology of RTT.

## Discussion

This study provides a comprehensive analysis of how PCB exposure influences cortical cell type-specific gene expression in *Mecp2e1* mosaic females, shedding light on the complex interplay between an environmental factor and an X-linked gene, each regulating neurodevelopment. Using a combination of single-nucleus RNA sequencing, differential expression analysis, and network analysis, we explored the effects of PCBs on RTT syndrome transcriptional networks and identified potential inter-cellular interactions. We were able to directly distinguish WT- and mutant -expressing cells within the HET mosaic brains, facilitating a detailed comparative analysis of gene expression and pathway dysregulation between WT- and mutant-expressing neurons within and across experimental genotype and treatment groups^7^. These experimental comparisons provided a high degree of specificity in parsing WT from mutant cells, which was crucial for understanding how PCB exposure impacts cell non-autonomous effects in the balance between inhibitory and excitatory neurons. The reduction in the number of RTT-associated dysregulated pathways observed with PCB exposure indicates a mitigating effect of PCB exposure on RTT-related gene dysregulation in both mouse model and human cortices. Together, these results provide multiple novel findings relevant to understanding the complex cellular and transcriptional pathogenesis of both Rett syndrome and PCB neurotoxicity.

Our prior analysis of Rett syndrome at the single cell transcriptome level revealed that in *HET* female mice the WT-expressing neurons were dynamically responsive to the mosaic HET cortical environment^7^. In the current study, both DEG and pathway analyses showed that PCB exposure led to fewer dysregulated genes compared to vehicle treatment in the RTT mutant mouse background. Parsing of the cells into mutant- and WT-expressing neurons demonstrated that PCB primarily impacted the WT-expressing cell transcriptome. This finding indicates a possible protective role of PCB exposure in the unusual mosaic context of Rett syndrome pathogenesis. The KEGG pathways commonly affected across all conditions included those relevant to adaptive pathways relevant to synaptic regulation, such as circadian rhythms and cAMP signaling, highlighting the broad impact of PCB exposure on critical homeostatic information networks between cells that are dysregulated in RTT.

High dimensional Weighted Gene Co-Expression Network Analysis (hdWGCNA) further elucidated the effects of PCB exposure and *Mecp2* mutation on gene networks^25^. We identified seven distinct gene modules, with multiple modules, including brown, that showed a significant positive correlation with *Mecp2* genotype in all cell types, but differed in the direction of correlations with PCB treatment. PCBs were negatively associated with brown module transcript levels predominantly in GABAergic neuronal subtypes. Interestingly, the female-specific long noncoding RNA *Xist* was a member of the brown network. Furthermore, there was a significant interaction between *Mecp2e1* genotype and PCB exposure that was cell type-dependent, with GABAergic neurons being the cell type with the strongest interaction effect. Somatostatin (Sst) cells, a GABAergic neuronal subtype, was found to be particularly strongly influenced by PCB exposure and genetic mutations. Together, these results suggest that *Xist* and other genes within the brown module may mediate the interactions between *Mecp2* genotype and PCB exposure in cortical GABAergic neurons. Future research could determine if interactions between PCBs and *Mecp2* are specific to females and if *Xist* is required for interaction effects in females.

While the study provides valuable insights, several limitations should be considered. Our analysis focused specifically on PCB exposure, but other environmental factors may also influence the inter-cellular transcriptional networks in the pathogenesis of Rett syndrome^26^. Future research should explore a broader range of environmental exposures impacting Rett syndrome severity to gain a more comprehensive understanding. Longitudinal studies are needed to assess how gene expression changes evolve over time and their long-term implications for RTT and if different PCB doses change to effects on transcriptional and disease phenotypes^27^. Additionally, further functional validation of the identified pathways and gene modules is necessary to confirm their roles in RTT and the impact of PCB exposure.

In conclusion, this study advances our understanding of how PCB exposure interacts with *Mecp2e1* mutations to influence RTT molecular pathogenesis in female mouse and human cortex. By elucidating the effects on gene expression, pathway dysregulation, and network dynamics, we could focus on therapeutic targets and mechanisms that could inform future research and clinical strategies for managing RTT.

## Acknowledgements

We would like to express our gratitude to the members of the Flow Cytometry Core, Bridget McLaughlin and Jonathan Van Dyke for their technical assistance and expertise in sorting nuclei. This research was supported by NIH T32ES007059, R01ES029213, R01AA027075, S10OD010786, U2CES030158, and U24DK097154.

## Author’s Contributions

OS, KEN and JML designed the study. JML and OS acquired funding for the study. KEN and AV performed the PCB dosing and dissections of the mice. OS and DHY performed the nuclei isolation and single cell assays. OS performed the bioinformatic analyses with intellectual contributions from IK. OS wrote the manuscript with intellectual contributions from KEN, PJL, DHY and JML. CGT and OS generated all the figures. All authors reviewed and approved the final manuscript.

## STAR Methods

### Resource Availability

#### Lead Contact

Further information and requests for resources and reagents should be directed to and will be fulfilled by the lead contact, Janine M. LaSalle.

## Materials Availability

This study did not generate new unique reagents.

## Data and Code Availability

- Raw and processed sequencing data has been deposited at GEO and is publicly available as of the date of publication.
- All original code has been deposited at GitHub and is publicly available as of the date of publication (https://github.com/osmansharifi/PCB_mouse_snRNAseq).
- Any additional information required to reanalyze the data reported in this paper is available from the lead contact upon request.

## Experimental Model and Subject Details Mouse dosing

*Mecp2e1^-/+^* female mice were crossed with wild-type C57BL6/J males (Jackson Labs strain 000664) to generate the *Mecp2e1^-/+^* progeny used in this study. This breeding strategy follows the established model^19^ producing females with mosaic expression of MeCP2-e1 due to random X-chromosome inactivation. A PCB mixture modeling the 12 most prevalent congeners found in maternal serum from the ASD-enriched MARBLES cohort was prepared according to previously established protocols^8,28^. This formulation contained varying proportions of PCB congeners: PCB 28 (48.2%), PCB 11 (24.3%), PCB 118 (4.9%), PCB 101 (4.5%), PCB 52 (4.5%), PCB 153 (3.1%), PCB 180 (2.8%), PCB 149 (2.1%), PCB 138 (1.7%), PCB 84 (1.5%), PCB 135 (1.3%), and PCB 95 (1.2%). Female C57BL/6J mice (The Jackson Laboratory), ∼10 weeks of age, were administered either 0.01 mg/kg/day of the PCB mixture delivered via diet (in peanut butter) or vehicle control (peanut oil in peanut butter) for about 6 weeks. On week ∼16, dams (n_WT PCB-exposed_ =4, n*_Mecp2-e1_* _PCB-exposed_ =4, n_WT vehicle control_ =4, n*_Mecp2-e1_* _vehicle_ =4) were euthanized. From a total of 16 dams, cerebral cortices were harvested and used for single nucleus preparations. All experimental procedures received approval from the University of California, Davis Institutional Animal Care and Use Committee (IACUC).

## Methods Details

### Single-nucleus RNA sequencing

Single nucleus suspensions were prepared from each mouse brain according to previous stablished protocols^7,29^. A 3.0 mm² section of cortex was excised from each mouse brain and minced using a scalpel and subsequently homogenized in 0.5 ml of nuclei lysis buffer containing RNAse inhibitor (Roche, Indianapolis, ID). The homogenate was transferred to a larger vessel with an additional 1.0 ml of nuclei lysis buffer, thoroughly mixed, and incubated on ice for 5 minutes. Nuclei were separated from the lysate by filtration through a 70 μm FlowMi cell strainer (Sp-Belart, Wayne, NJ).

The nuclei were then pelleted by centrifugation at 500×G for 5 minutes at 4°C and resuspended in 1.5 ml of nuclei wash buffer for a 5-minute incubation. Following this incubation, nuclei were pelleted again under identical centrifugation conditions and washed twice in nuclei wash and resuspension buffer. The suspension was filtered through a 35 μm FlowMi filter (Sp-Belart, Wayne, NJ) and resuspended in nuclei wash and resuspension buffer containing 5 μg/ml DAPI. Nuclear concentration and integrity were assessed using a Countess cell counter (Fisher Scientific, Waltham, MA).

To remove debris and nuclear aggregates, the samples were processed on a MoFlow Astrios cell sorter (Beckman-Coulter, Brea, CA). Approximately 150,000 nuclei per sample were sorted and maintained on ice prior to library generation.

Single Cell 5′ Library V1 & Gel Bead Kits (10× Genomics, Pleasanton, CA) were employed for cDNA preparation and generation of barcoded and indexed snRNA-seq 5′ libraries following the manufacturer’s protocol. A target of 10,000 nuclei per sample was established. The snRNA-seq 5′ libraries were quantified using a Kapa library quantification kit (Roche, Indianapolis, IN) and subsequently pooled for sequencing. Paired-end sequences of 150 base pairs were generated using a NovaSeq S4 sequencer (Illumina, San Diego, CA). The mouse cortical samples yielded approximately 75,000 reads per cell and an average of 240,437,728 reads per sample.

### Quantification and Statistical Analyses Bioinformatic Analyses

Raw reads from mouse samples were aligned to the mm10-1.2.0 reference genome using Cellranger v.3.2.3. The resulting cell by gene count matrices were imported into R 4.2.2 to create Seurat objects using Seurat_4.3.0.1. Quality control filtering was applied to mouse samples using the following criteria: cells were required to have less than 7% mitochondrial content, between 200 and 5,625 expressed genes, and between 208 and 16,300 unique molecular identifiers (UMI). The expression counts were log-transformed, normalized, and scaled using Seurat 4.3.0.1. Sample metadata was enriched with information regarding genotype, treatment, and *Mecp2-e1* allele status. For cell type annotation, previously published single nucleus data from Rett mouse cortex served as a reference^7^. Cell type labels were transferred to the current dataset using scTransform for alignment. Cell marker testing was performed to validate the accuracy of cell type annotations. Visualization tools including UMAP plots and dotplots demonstrating validation of cell type-specific markers were generated using scCustomize 2.1.1^30^.

LimmaVoom was employed to identify differentially expressed genes (DEGs) across 15 distinct experimental comparisons. Prior to differential expression analysis, cell numbers were normalized by downsampling to match the condition with the fewest cells within each experiment. Complete parameters for all differential expression analyses are available in the GitHub repository. DEGs with a p-value ≤0.05 from each experiment were subsequently used as input for KEGG pathway analysis, performed using the enrichR 3.2 R package.

For hdWGCNA analysis, cells from both MARBLES mix-treated and vehicle conditions in the processed Seurat object were utilized as input. Additional covariates included pregnancy status, weight at euthanasia, and treatment exposure duration. Genes were required to be expressed in at least 5% of cells to be included in the analysis. A signed network with a softpower of 0.8 was employed, resulting in the identification of 7 modules. Module scores were computed using the UCell method. Standard pipelines from hdWGCNA 0.2.4 were followed, with detailed parameters available in the GitHub repository.

Parsing *Mecp2e1* alleles in the mosaic cortices was performed based on a previously published method^31^. *Mecp2* reads were extracted from the raw fastq files generated for each sample. abBLAST 3.0 and BWA 0.7.17 mem were used in combination to extract Mecp2 reads (via alleler.py). The reference for alignment consisted of 100 bp of the *Mecp2* gene, comprising 50 bp upstream and 50 bp downstream of the exon1 start codon. Within the aligned reads, *Mecp2e1* mutant (TTG) and wild-type (ATG) start codons were enumerated using alleler.py. Each read contained cell barcode and UMI information, which enabled the incorporation of mutant and wild-type cell information into the Seurat object metadata.

For differential expression analysis, each gene in each sample was modeled using a negative binomial distribution. The mean was calculated as the product of library size and relative abundance (gene expression levels), with variance as a function of the mean. Significant DEGs were identified based on an adjusted p-value ≤0.05, employing the Benjamini-Hochberg method to control for false discovery. These significant DEGs served as input for KEGG pathway analysis, which utilized Fisher’s exact test to determine if the overlap between input DEGs and background genes was significant (adjusted p-value ≤0.05). The hdWGCNA analysis utilized Pearson correlation tests to compare gene expression with phenotypic data, with significant correlations determined based on an adjusted p-value ≤0.05.

---

**Supplemental Figure 1.** : Bar graph showing proportion of cell types across the four experimental groups.

**Supplemental Figure 2.** : Bioinformatic pipeline for parsing out *Mecp2e1* WT- and mutant-expressing cells in the mosaic RTT cortex and differential analysis

**Supplemental Figure 3.** : Heatmap showing the correlations between gene expression and phenotypes. In the context of genotype, a positive correlation indicates increased transcript levels in RTT mice, while a positive correlation with treatment indicated increased transcript levels with PCBs. Conversely, a positive correlation with pregnancy means the mice are not pregnant, whereas a negative correlation indicates that the mice are pregnant. * adjusted-p-value< 0.05, **adjusted-p- value < 0.01, ***adjusted p-value <0.001

**Supplemental Figure 4.** Interaction plot of *Xist* expression with *Mecp2e1* mutant allele, treatment and cell type.

**Supplemental Table 1.** :Cellular proportions per cell type per experimental condition.

**Supplemental Table 2.** List of significant KEGG terms per DEG test for each cell type.

**Supplemental Table 3.** Number of significant DEGs per DEG experiment for GABAergic, glutamatergic and non-neuronal cells.

**Supplemental Table 4.** List of significant DEGs per cell type for 15 different DEG experiments.

## References

1. Amir, R.E., Veyver, I.B.V.D., Wan, M., Tran, C.Q., Francke, U., and Zoghbi, H.Y. (1999). Rett syndrome is caused by mutations in X-linked MECP2, encoding methyl-CpG-binding protein 2. Nat. Genet. 23, 185–188. 10.1038/13810.

2. Zoghbi, H.Y. (2005). MeCP2 Dysfunction in Humans and Mice. J. Child Neurol. 10.1177/08830738050200090701.

3. Chahrour, M., Sung, Y.J., Shaw, C., Zhou, X., Wong, S.T.C., Qin, J., and Zoghbi, H.Y. (2008). MeCP2, a key contributor to neurological disease, activates and represses transcription. Science. 10.1126/science.1153252.

4. Jordan, C., Li, H.H., Kwan, H.C., and Francke, U. (2007). Cerebellar gene expression profiles of mouse models for Rett syndrome reveal novel MeCP2 targets. BMC Med. Genet. 8, 36. 10.1186/1471-2350-8-36.

5. Tudor, M., Akbarian, S., Chen, R.Z., and Jaenisch, R. (2002). Transcriptional profiling of a mouse model for Rett syndrome reveals subtle transcriptional changes in the brain. Proc. Natl. Acad. Sci. U. S. A. 99. 10.1073/pnas.242566899.

6. Neul, J.L., Fang, P., Barrish, J., Lane, J., Caeg, E.B., Smith, E.O., Zoghbi, H., Percy, A., and Glaze, D.G. (2008). Specific mutations in methyl-CpG-binding protein 2 confer different severity in Rett syndrome. Neurology 70, 1313–1321. 10.1212/01.wnl.0000291011.54508.aa.

7. Sharifi, O., Haghani, V., Neier, K.E., Fraga, K.J., Korf, I., Hakam, S.M., Quon, G., Johansen, N., Yasui, D.H., and LaSalle, J.M. (2024). Sex-specific single cell-level transcriptomic signatures of Rett syndrome disease progression. Commun. Biol. 7, 1292. 10.1038/s42003-024-06990-0.

8. Laufer, B.I., Neier, K., Valenzuela, A.E., Yasui, D.H., Schmidt, R.J., Lein, P.J., and LaSalle, J.M. (2022). Placenta and fetal brain share a neurodevelopmental disorder DNA methylation profile in a mouse model of prenatal PCB exposure. Cell Rep. 38, 110442. 10.1016/j.celrep.2022.110442.

9. Park, H.-Y., Hertz-Picciotto, I., Sovcikova, E., Kocan, A., Drobna, B., and Trnovec, T. (2010). Neurodevelopmental toxicity of prenatal polychlorinated biphenyls (PCBs) by chemical structure and activity: a birth cohort study. Environ. Health 9, 51. 10.1186/1476-069X-9-51.

10. Berghuis, S.A., Bos, A.F., Sauer, P.J.J., and Roze, E. (2015). Developmental neurotoxicity of persistent organic pollutants: an update on childhood outcome. Arch. Toxicol. 10.1007/s00204-015-1463-3.

11. Jacobson, J.L., and Jacobson, S.W. (2003). Prenatal exposure to polychlorinated biphenyls and attention at school age. J. Pediatr. 143, 780–788. 10.1067/S0022-3476(03)00577-8.

12. Lesiak, A., Zhu, M., Chen, H., Appleyard, S.M., Impey, S., Lein, P.J., and Wayman, G.A. (2014). The environmental neurotoxicant PCB 95 promotes synaptogenesis via ryanodine receptor-dependent miR132 upregulation. J. Neurosci. 10.1523/JNEUROSCI.2884-13.2014.

13. Wayman, G.A., Yang, D., Bose, D.D., Lesiak, A., Ledoux, V., Bruun, D., Pessah, I.N., and Lein, P.J. (2012). PCB-95 Promotes Dendritic Growth via Ryanodine Receptor–Dependent Mechanisms. Environ. Health Perspect. 120, 997–1002. 10.1289/ehp.1104832.

14. Mitchell, M.M., Woods, R., Chi, L.H., Schmidt, R.J., Pessah, I.N., Kostyniak, P.J., and Lasalle, J.M. (2012). Levels of select PCB and PBDE congeners in human postmortem brain reveal possible environmental involvement in 15q11-q13 duplication autism spectrum disorder. Environ. Mol. Mutagen. 10.1002/em.21722.

15. Dunaway, K.W., Islam, M.S., Coulson, R.L., Lopez, S.J., Vogel Ciernia, A., Chu, R.G., Yasui, D.H., Pessah, I.N., Lott, P., Mordaunt, C., et al. (2016). Cumulative Impact of Polychlorinated Biphenyl and Large Chromosomal Duplications on DNA Methylation, Chromatin, and Expression of Autism Candidate Genes. Cell Rep. 10.1016/j.celrep.2016.11.058.

16. Granillo, L., Sethi, S., Keil, K.P., Lin, Y., Ozonoff, S., Iosif, A.-M., Puschner, B., and Schmidt, R.J. (2019). Polychlorinated biphenyls influence on autism spectrum disorder risk in the MARBLES cohort. Environ. Res. 171, 177–184. 10.1016/j.envres.2018.12.061.

17. Keil Stietz, K.P., Sethi, S., Klocke, C.R., de Ruyter, T.E., Wilson, M.D., Pessah, I.N., and Lein, P.J. (2021). Sex and Genotype Modulate the Dendritic Effects of Developmental Exposure to a Human-Relevant Polychlorinated Biphenyls Mixture in the Juvenile Mouse. Front. Neurosci. 15, 766802. 10.3389/fnins.2021.766802.

18. Sethi, S., Keil Stietz, K.P., Valenzuela, A.E., Klocke, C.R., Silverman, J.L., Puschner, B., Pessah, I.N., and Lein, P.J. (2021). Developmental Exposure to a Human-Relevant Polychlorinated Biphenyl Mixture Causes Behavioral Phenotypes That Vary by Sex and Genotype in Juvenile Mice Expressing Human Mutations That Modulate Neuronal Calcium. Front. Neurosci. 15, 766826. 10.3389/fnins.2021.766826.

19. Yasui, D.H., Gonzales, M.L., Aflatooni, J.O., Crary, F.K., Hu, D.J., Gavino, B.J., Golub, M.S., Vincent, J.B., Schanen, N.C., Olson, C.O., et al. (2014). Mice with an isoform-ablating Mecp2exon 1 mutation recapitulate the neurologic deficits of Rett syndrome. Hum. Mol. Genet. 10.1093/hmg/ddt640.

20. Tasic, B., Yao, Z., Graybuck, Lucas, Smith, K.A., Nghi Nguyen, Thuc, Bertagnolli, D., Goldy, J., Garren, Emma, Cconomo, M.N., Viswanathan, S., et al. (2018). Shared and distinct transcriptomic cell types across neocortical areas. Nature. 10.1038/s41586-018-0654-5.

21. Bullert, A.J., Wang, H., Valenzuela, A.E., Neier, K., Wilson, R.J., Badley, J.R., LaSalle, J.M., Hu, X., Lein, P.J., and Lehmler, H.-J. (2024). Interactions of Polychlorinated Biphenyls and Their Metabolites with the Brain and Liver Transcriptome of Female Mice. ACS Chem. Neurosci. 15, 3991–4009. 10.1021/acschemneuro.4c00367.

22. Medehouenou, T.C.M., Ayotte, P., Carmichael, P.-H., Kröger, E., Verreault, R., Lindsay, J., Dewailly, É., Tyas, S.L., Bureau, A., and Laurin, D. (2019). Exposure to polychlorinated biphenyls and organochlorine pesticides and risk of dementia, Alzheimer’s disease and cognitive decline in an older population: a prospective analysis from the Canadian Study of Health and Aging. Environ. Health 18, 57. 10.1186/s12940-019-0494-2.

23. Hatcher-Martin, J.M., Gearing, M., Steenland, K., Levey, A.I., Miller, G.W., and Pennell, K.D. (2012). Association between polychlorinated biphenyls and Parkinson’s disease neuropathology. NeuroToxicology 33, 1298–1304. 10.1016/j.neuro.2012.08.002.

24. Bakken, T.E., Jorstad, N.L., Hu, Q., Lake, B.B., Tian, W., Kalmbach, B.E., Crow, M., Hodge, R.D., Krienen, F.M., Sorensen, S.A., et al. Comparative cellular analysis of motor cortex in human, marmoset and mouse. 10.1038/s41586-021-03465-8.

25. Morabito, S., Reese, F., Rahimzadeh, N., Miyoshi, E., and Swarup, V. (2023). hdWGCNA identifies co-expression networks in high-dimensional transcriptomics data. Cell Rep. Methods 3. 10.1016/j.crmeth.2023.100498.

26. Nag, N., Ward, B., and Berger-Sweeney, J.E. (2009). Nutritional factors in a mouse model of Rett syndrome. Neurosci. Biobehav. Rev. 33, 586–592. 10.1016/j.neubiorev.2008.03.007.

27. Tarquinio, D.C., Hou, W., Berg, A., Kaufmann, W.E., Lane, J.B., Skinner, S.A., Motil, K.J., Neul, J.L., Percy, A.K., and Glaze, D.G. (2017). Longitudinal course of epilepsy in Rett syndrome and related disorders. Brain 140, 306–318. 10.1093/brain/aww302.

28. Sethi, S., Morgan, R.K., Feng, W., Lin, Y., Li, X., Luna, C., Koch, M., Bansal, R., Duffel, M.W., Puschner, B., et al. (2019). Comparative Analyses of the 12 Most Abundant PCB Congeners Detected in Human Maternal Serum for Activity at the Thyroid Hormone Receptor and Ryanodine Receptor. Environ. Sci. Technol. 53, 3948–3958. 10.1021/acs.est.9b00535.

29. Martelotto, L. (2018). ‘ Frankenstein ’ protocol for nuclei isolation from fresh and frozen tissue followed by 10x Genomics (Luciano Martelotto). 3–4.

30. Samuel Marsh, Maëlle Salmon, Paul Hoffman, and kew24 (2024). samuel-marsh/scCustomize: Version 3.0.1. Version v3.0.1 (Zenodo). 10.5281/ZENODO.5706430.

31. Sharifi, O., Haghani, V., Fraga, K., Korf, I., Neier, K., and Lyman, H. (2024). osmansharifi/snRNA-seq-pipeline: Rett snRNA-seq cortex. Version Release-1.02 (Zenodo). 10.5281/zenodo.13761244.

