## Supplementary figures and images for "Female cortical cellular mosaicism underlies shared MeCP2 and PCB impacted gene pathways"

### Supplemental figure 1

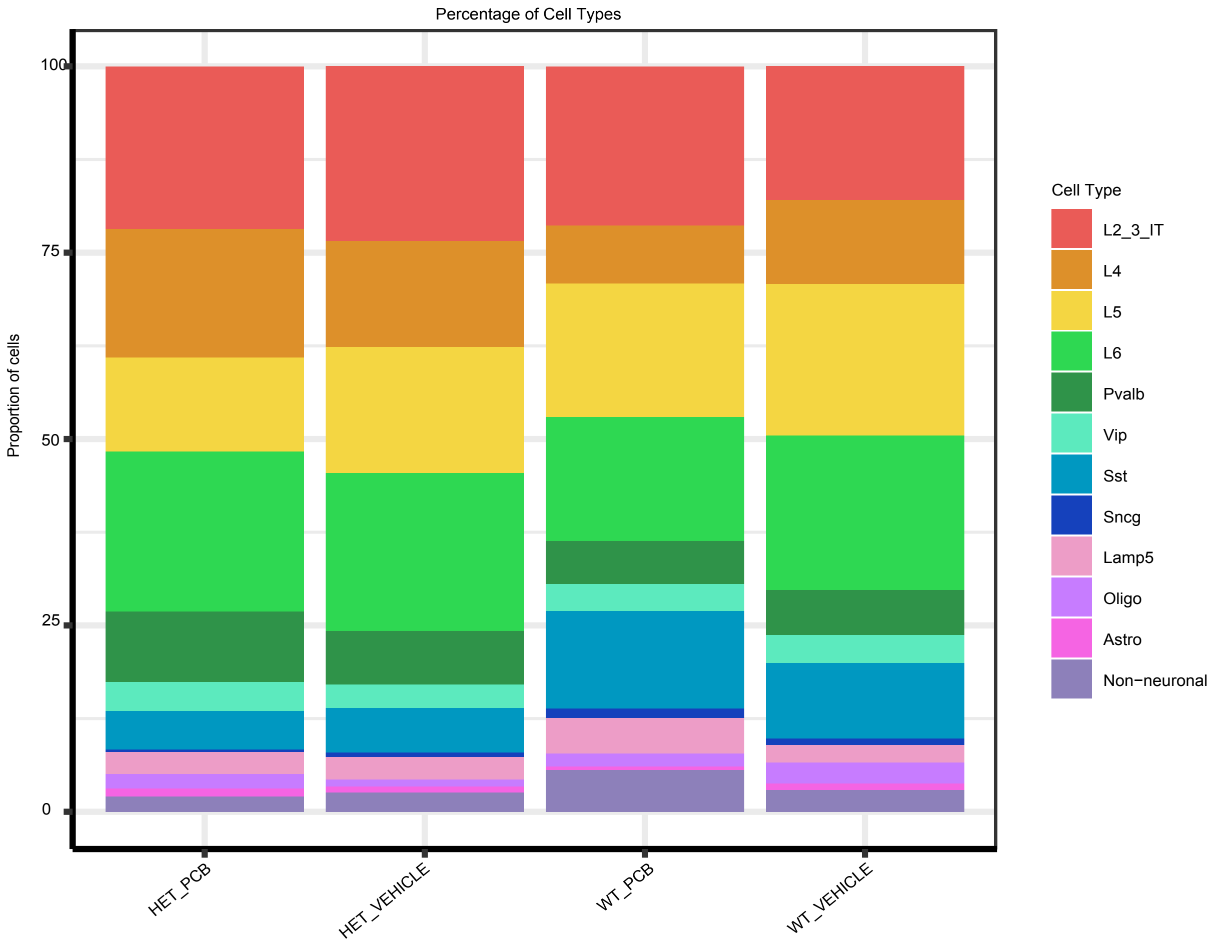

### Supplemental figure 2

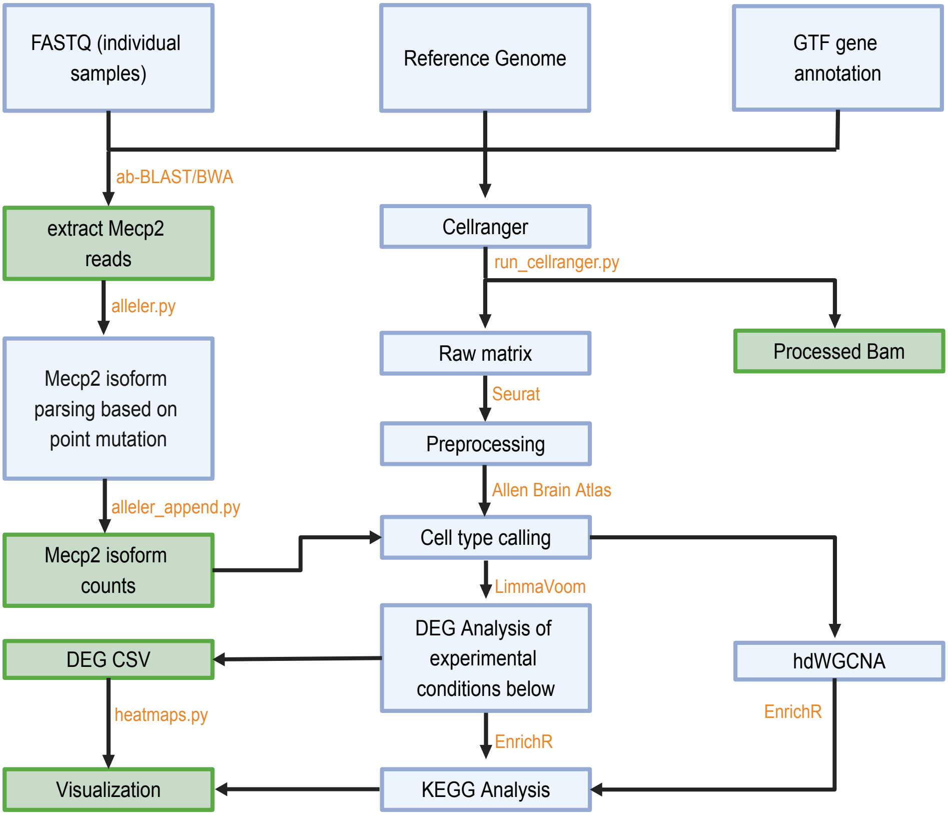

### Supplemental figure 3

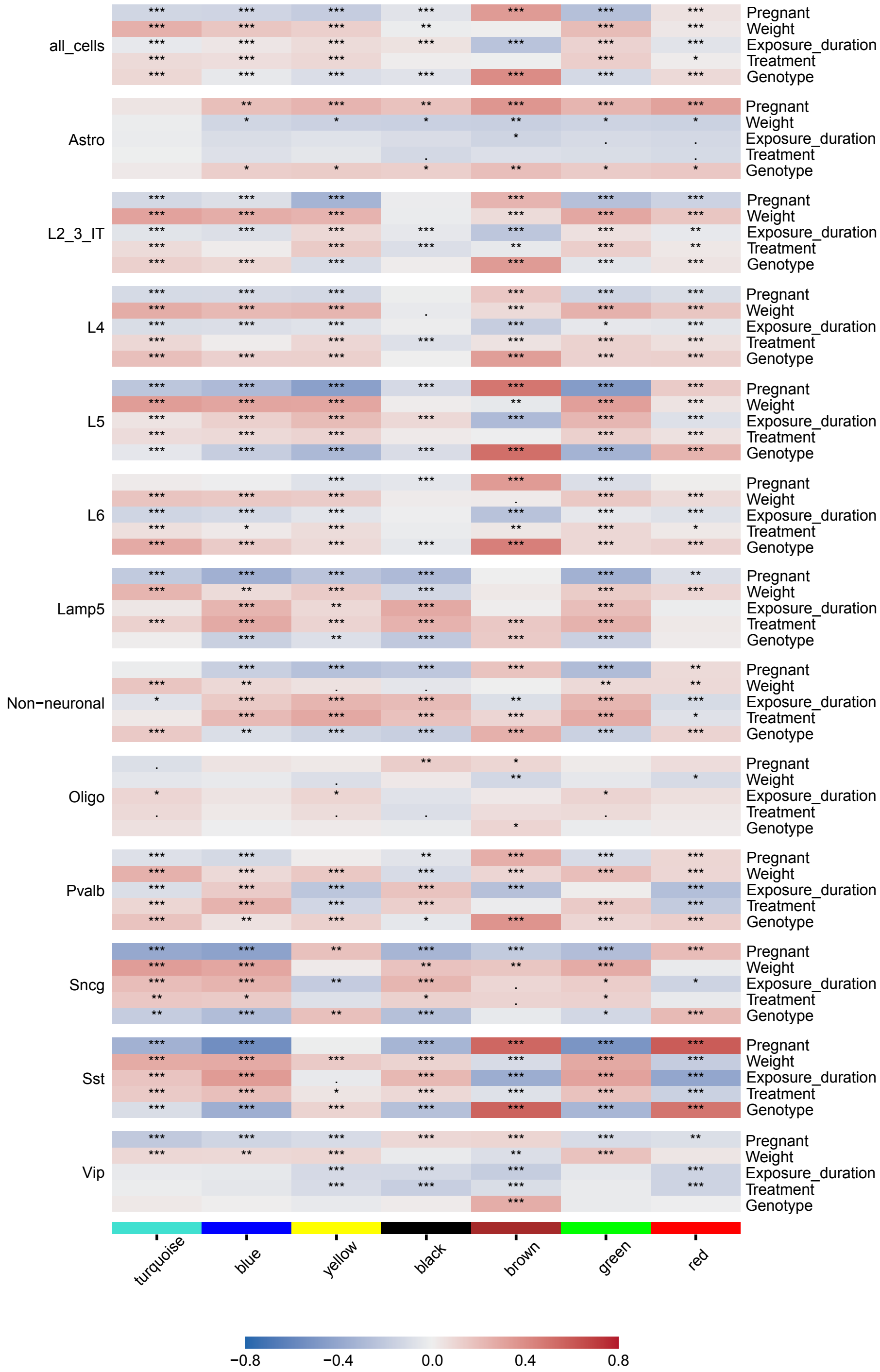
