## Supplemental figure 4 for "Female cortical cellular mosaicism underlies shared MeCP2 and PCB impacted gene pathways"

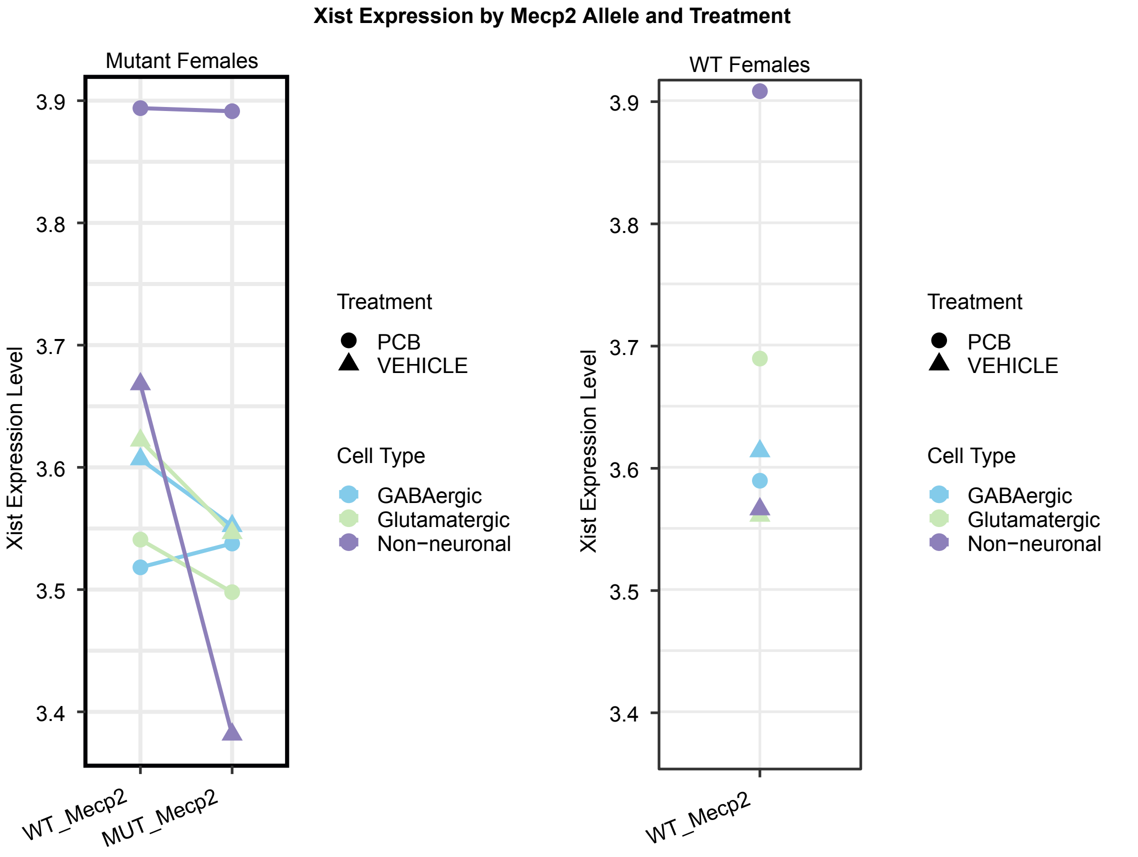


| 3-way ANOVA | Df | Sum Sq | Mean Sq | F value | Pr(>F) | Significance |
|---|---|---|---|---|---|---|
| Mecp2_allele | 1 | 1.2 | 1.2216 | 3.608 | 0.0577 | . |
| Treatment | 1 | 0.5 | 0.538 | 1.589 | 0.2077 | |
| Cell Type | 2 | 0.3 | 0.1534 | 0.453 | 0.6357 | |
| Mecp2_allele:Treatment | 1 | 0.2 | 0.1967 | 0.581 | 0.446 | |
| Mecp2_allele:Cell Type | 2 | 0.3 | 0.1346 | 0.397 | 0.6721 | |
| Treatment:Cell Type | 2 | 2.3 | 1.1258 | 3.325 | 0.0363 | * |
| Mecp2_allele:Treatment:Cell Type | 2 | 0.2 | 0.0762 | 0.225 | 0.7986 | |
| Residuals | 1370 | 463.8 | 0.3386 | | | |
| --- | | | | | | |
| Signif. codes: | 0 ‘***’ | 0.001 ‘**’ | 0.01 ‘*’ | 0.05 ‘.’ | 0.1 ' ' | 1 |
